# The HosA histone deacetylase regulates stress resistance, host cell interactions, and virulence in *Aspergillus fumigatus*

**DOI:** 10.1101/2025.11.19.689210

**Authors:** Hong Liu, Pamela Lee, Alice Vo, Sanjoy Paul, Quynh T. Phan, Jianfeng Lin, Vincent M. Bruno, Mark A. Stamnes, W. Scott Moye-Rowley, Scott G. Filler

## Abstract

The capacity of *Aspergillus fumigatus* to cause invasive pulmonary aspergillosis depends on its ability to adapt to dynamic and stressful microenvironments within the host. Epigenetic regulation, including histone deacetylation, plays a critical role in fungal adaptation to stress. Here, we investigated the role of the class I histone deacetylase (HDAC) HosA in *A. fumigatus* stress resistance, host cell interactions, and virulence. A Δ*hosA* mutant had increased susceptibility to intracellular oxidant stress induced by menadione. It also had impaired capacity to invade and damage two pulmonary epithelial cell lines *in vitro*. In a corticosteroid-immunosuppressed mouse model of invasive aspergillosis, mice infected with the Δ*hosA* mutant survived significantly longer than those infected with the wild-type strain, despite having similar pulmonary fungal burden. The Δ*hosA* mutant also induced a weaker inflammatory response than the wild-type strain. Transcriptomic analysis revealed that HosA regulates genes involved in secondary metabolite biosynthesis and energy metabolism, functioning as both an activator and repressor of distinct gene sets. Collectively, these results indicate that HosA is a key epigenetic regulator that governs *A. fumigatus* interactions with host cells and virulence during invasive pulmonary aspergillosis.

**Importance:** Epigenetic modifications in *A. fumigatus* can be induced by environmental changes and stresses such as those induced by interaction with host cells. HosA, a class I histone deacetylase, has been shown to play a key role in regulating secondary metabolism in several *Aspergillus* species, but its function in *A. fumigatus* was previously unknown. We found that deletion of *hosA* increased susceptibility to intracellular, but not extracellular, oxidative stress. The Δ*hosA* mutant also exhibited significantly reduced pulmonary epithelial cell invasion and host cell damage, as well as attenuated virulence in immunosuppressed mice. Together, these findings indicate that HosA functions as a key epigenetic regulator that governs stress resistance, secondary metabolism, and fungal-host interactions. Defining the functions of HosA could provide critical insight into the epigenetic mechanisms that control fungal pathogenicity and may reveal a potential therapeutic target for invasive aspergillosis.

## Introduction

The capacity of *Aspergillus fumigatus* to cause invasive pulmonary aspergillosis is due to its ability to invade the epithelial lining of the airways and then survive and proliferate within the lung parenchyma. During these processes, the fungus is exposed to a variety of stressors induced by the host. These stressors include hypoxia, attack by phagocytic cells, and limitation of essential nutrients such as nitrogen, iron, and zinc (1–5). The ability of *A. fumigatus* to rapidly respond to stresses in the various microenvironments of the host is essential for the fungus to proliferate and cause disease. The response of *A. fumigatus* to stressors is governed by transcription factors and protein kinases (6–8). It is also regulated by epigenetic factors, such as histone modification.

One key class of epigenetic regulators are histone deacetylases (HDACs). These enzymes remove acetyl groups from the *ε*-amino group of lysine residues in histones and other proteins. HDACs can both inhibit and stimulate gene expression. Based on homology to HDACs in *Aspergillus niger* and *Aspergillus nidulans* (9, 10)*, A. fumigatus* possesses three classes of HDACs, comprising a total of seven different proteins. The two class I HDACs are HosA (Afu2g03810) and RpdA (Afu2g03390). RpdA was reported to essential in *A. fumigatus* (11). Although the function of HosA in *A. fumigatus* has not been studied previously, it has been found to be required for normal growth, pigmentation, and conidiation in *A. niger, A. nidulans* and *Aspergillus flavus* (9, 12, 13). In these fungi, HosA also governs the production of secondary metabolites.

Here we investigated the function of HosA in *A. fumigatus* pathogenicity. We found that a Δ*hosA* mutant had slightly reduced growth rate on minimal medium, impaired interactions with pulmonary epithelial cells in vitro, and attenuated virulence in the immunosuppressed mouse model of invasive pulmonary aspergillosis.

## Results

### HosA mediates resistance to intracellular oxidative stress

To study the function of HosA in *A. fumigatus*, a mutant deleted for *hosA* (Afu2g03810) was constructed using CRISPR/cas9. The resulting Δ*hosA* mutant had near normal radial growth on nutrient-rich Sabouraud dextrose agar, although the edges of the colony were irregular, in contrast to smooth edges of colonies of the wild-type parent and Δ*hosA*+*hosA* complemented strain (Fig. 1A). Under this growth condition, the Δ*hosA* mutant also had a defect in conidiation, producing 33% fewer conidia than the wild-type strain (Fig. 1B) On the relatively nutrient-poor *Aspergillus* minimal medium, the Δ*hosA* mutant grew slightly slower than the wild-type strain (Fig. 1C). Thus, HosA plays a modest role in controlling conidiation and growth.

**Fig. 1.**
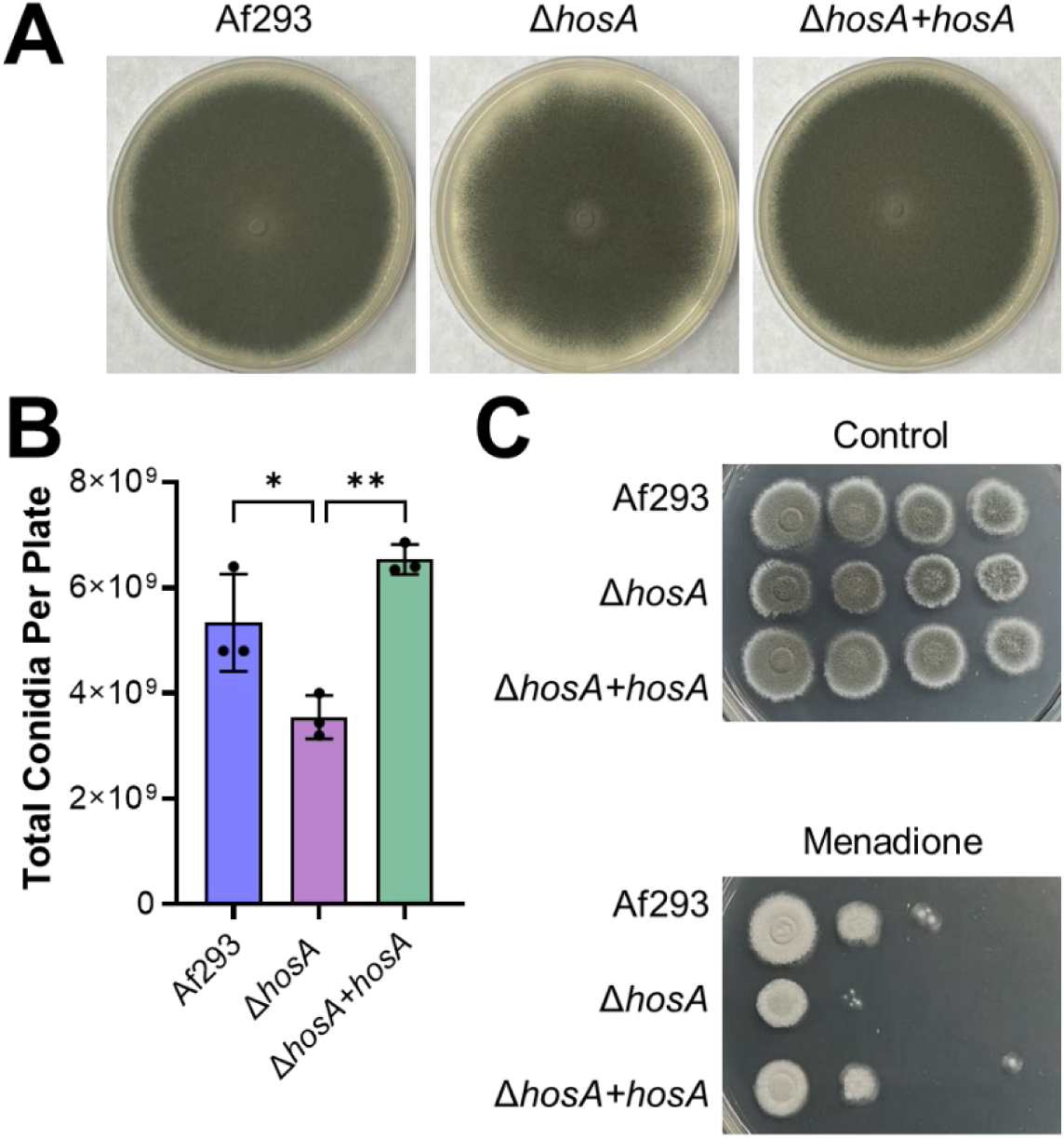
The Δ*hosA* mutant had reduced conidiation and increased susceptibility to intracellular oxidative stress. (A) Radial growth of the wild-type (Af293), Δ*hosA,* and Δ*hosA +hosA A. fumigatus* strains. Colonies were imaged after growth at 37°C for 5 days. (B) Conidia production by the indicated *A. fumigatus* strains after growth on Sabouraud dextrose agar in a 100 mm petri dish at 37°C for 5 days. Results are mean ± SD of 3 independent experiments. (C) Susceptibility of the indicated strains to 15 µM menadione. Plates were imaged after growth at 37°C for 48 h. *, *p* < 0.05; **, *p* < 0.01 by one-way ANOVA with Šídák’s multiple comparisons test.

Next, we investigated the susceptibility of Δ*hosA* mutant to various stressors. This mutant was more susceptible than the wild-type strain to intracellular oxidant stress induced by menadione, and this defect was rescued in the Δ*hosA*+*hosA* complemented strain. However, the Δ*hosA* mutant grew similarly to the wild-type strain in the presence of H_2_O_2_, indicating that it had increased susceptibility to intracellular, but not extracellular oxidant stress (Fig. S1A). The Δ*hosA* mutant also had similar susceptibility as the wild-type strain to Congo red, calcafluor white, protamine sulfate, voriconazole, and hypoxia (Fig. S1). These results indicate that HosA is required for resistance to intracellular, but not extracellular oxidant stress.

### HosA governs *A. fumigatus* invasion and damage of pulmonary epithelial cells

The capacity of *A. fumigatus* to invade and damage pulmonary epithelial cells *in vitro* is strongly associated with virulence in mouse models of infection (14–17). We investigated the role of HosA in governing the pathogenic interactions of *A. fumigatus* germlings with pulmonary epithelial cells. We used two human pulmonary epithelial cell lines, A549 cells, which are type II-like alveolar epithelial cells and HSAEC1-KT human small airway epithelial (HSAE) cells. We found that deletion of *hosA* had no significant effect on the number of germlings that were cell-associated with either epithelial cell line, indicating that HosA does not govern adherence to these cells (Fig. 2A and B). In contrast, the endocytosis of the Δ*hosA* germlings by the two epithelial cell lines was significantly reduced (Fig. 2C and D). The Δ*hosA* mutant also caused significantly less damage to both types of host cells. The epithelial cell invasion and damage defects of the Δ*hosA* mutant were rescued by integration of an intact copy of *hosA*. Collectively, these results indicate that HosA plays a key role in controlling the capacity of *A. fumigatus* to invade and damage pulmonary epithelial cells *in vitro*.

**Fig. 2.**
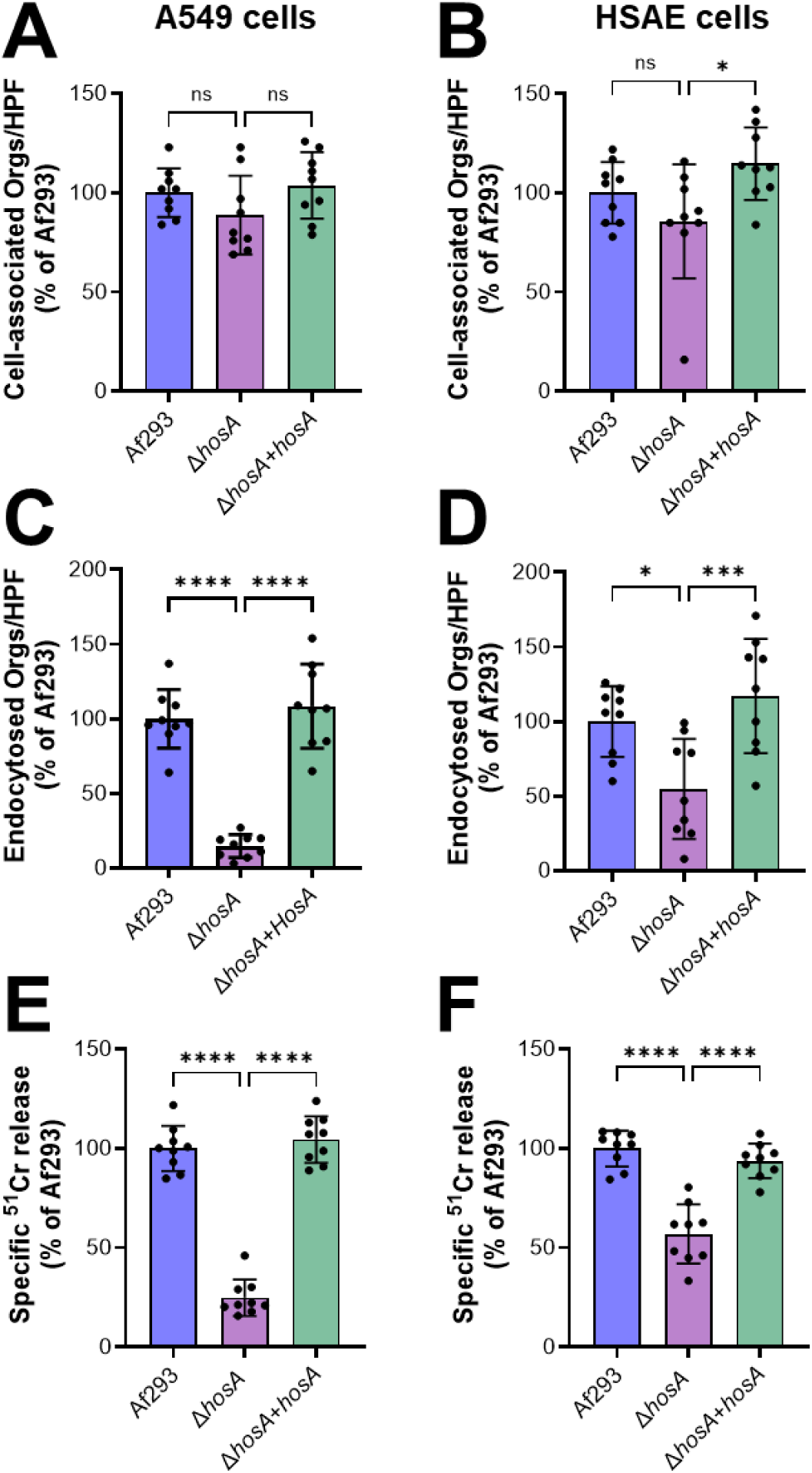
Reduced pulmonary epithelial cell invasion and damage of the Δ*hosA* mutant. (A-D). Cell-association (A and B) and endocytosis (C and D) of germlings of the indicated *A. fumigatus* strains by the A549 alveolar epithelial cell line (A and C) and the HSAE-KT human small airway epithelial (HSAE) cell line after a 2.5-h incubation. (E and F). Damage to A549 and HSAE cells by the indicated *A. fumigatus* strains after an 18-h incubation. Results are mean ± SD of 3 independent experiments, each performed in triplicate. ns, not significant; *, *p* < 0.05; ***, *p* < 0.001, ****, *p* < 0.0001 by one way ANVO with Šídák’s multiple comparisons test.

### HosA controls lethality independently of lung fungal burden in the mouse model of invasive pulmonary aspergillosis

To determine whether HosA governs *A. fumigatus* virulence, we used a mouse model of invasive pulmonary aspergillosis in which the animals were immunosuppressed with triamcinolone (18, 19). Mice infected with the Δ*hosA* mutant survived significantly longer than mice infected with either the wild-type parent strain or the Δ*hosA*+*hosA* complemented strain (Fig. 3A). By contrast, the pulmonary fungal burden of mice infected with the Δ*hosA* mutant was similar to that of mice infected with the wild-type strain (Fig. 3B). These results suggested that HosA governs disease outcome independently of fungal proliferation in the lung. The Δ*hosA* mutant also induced a weaker inflammatory response. Mice infected with this mutant had significantly lower levels of IL-1α, IL-6, CCL2, CXCL1, and CXCL2 in their lungs relative to mice infected with the wild-type strains, while the levels of CXCL5 and GM-CSF were unchanged (Fig. 3C). These results suggest that the Δ*hosA* mutant induced a weak host inflammatory response, which led to decreased lethality even though the lung fungal burden remained high.

**Fig. 3.**
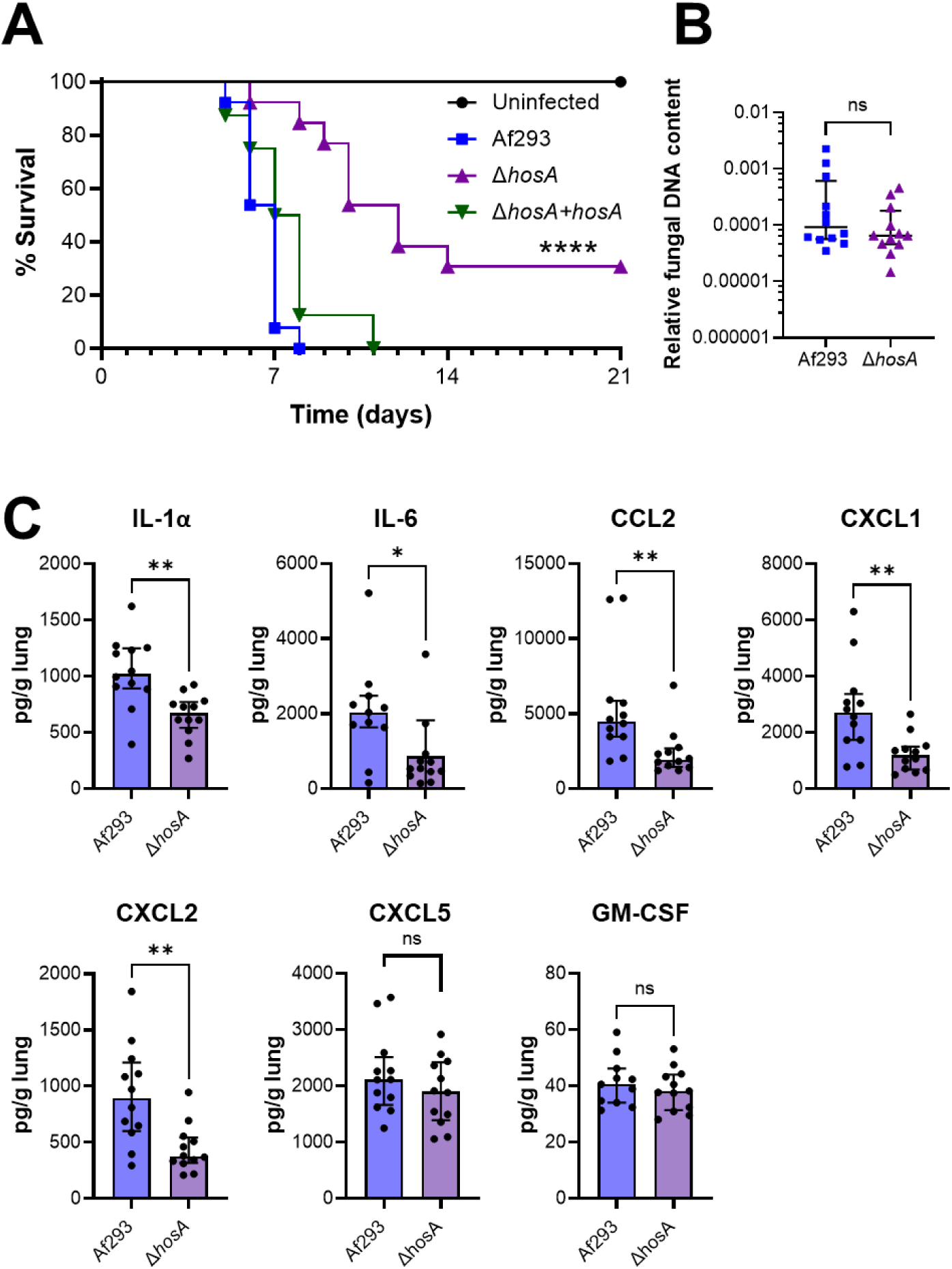
Attenuated virulence of the Δ*hosA* mutant. (A) Survival of corticosteroid immunosuppressed mice after pulmonary infection with the indicated *A. fumigatus* strains. Data are the combined results of 2 independent experiments for a total of 16 mice per strain of *A. fumigatus*. A total of 10 control mice were immunosuppressed, but not infected. (B) Lung fungal burden as determined by qPCR for fungal DNA after 5 d of infection. (C) Whole lung levels of the indicated cytokines, measured after 5 days of infection. Data in (C and D) are median ± interquartile range of a single experiment with 12 mice per strain. ns, not significant; *, *p* < 0.05; **, *p* < 0.01, ****, *p* < 0.0001 by the logrank test (A) or the Mann Whitney test (B and C).

### HosA governs expression of secondary metabolite and cellular energetic genes

We used RNA-seq to analyze the effects of *hosA* deletion on the *A. fumigatus* transcriptome. Organisms were grown for 24 h in Aspergillus minimal medium containing low zinc and iron, a condition that we found previously induces a transcriptional response that is similar to that elicited by growth in the lungs of immunosuppressed mice (6). In the Δ*hosA* mutant, 495 genes were significantly down-regulated and 377 genes were significantly up-regulated as compared to the wild-type strain (absolute log_2_ fold-change >1, adjusted p-value <0.05) (Table S1). Gene ontology (GO) term analysis of the down-regulated genes revealed enrichment for secondary metabolite biosynthesis, including fumitremorgin B and fumigermin (Table 1). GO term analysis of the up-regulated genes showed enrichment for genes involved in ATP production and the synthesis of fumiquinazoline C (Table 2). Thus, HosA is both a positive and negative regulator of secondary metabolite genes.

**Table 1.** Gene ontology (GO) term analysis of the genes that were significantly down-regulated in the Δ*hosA* mutant as compared to the wild-type strain.

| ID | Name | Odds ratio | P-value (Bonferroni adjusted) |
| --- | --- | --- | --- |
| GO:0044550 | Secondary metabolite biosynthetic process | 3.47 | $1.6 \times 10^{-10}$ |
| GO:0019748 | Secondary metabolic process | 3.3 | $1.1 \times 10^{-9}$ |
| GO:0055085 | Transmembrane transport | 2.52 | $1.9 \times 10^{-7}$ |
| GO:1900772 | Fumitremorgin B biosynthetic process | 67.21 | $2.4 \times 10^{-5}$ |
| GO:1900770 | Fumitremorgin B metabolic process | 67.21 | $2.4 \times 10^{-5}$ |
| GO:0009820 | Alkaloid metabolic process | 8.5 | 0.00081 |
| GO:0035835 | Indole alkaloid biosynthetic process | 8.76 | 0.0020 |
| GO:0035834 | Indole alkaloid metabolic process | 8.76 | 0.0020 |
| GO:0009821 | Alkaloid biosynthetic process | 8.03 | 0.0038 |
| GO:0140446 | Fumigermin biosynthetic process | 76.35 | 0.028 |
| GO:0008643 | Carbohydrate transport | 4.33 | 0.041 |

**Table 2.** Gene ontology (GO) term analysis of the genes that were significantly up-regulated in the Δ*hosA* mutant as compared to the wild-type strain.

| ID | Name | Odds ratio | P-value (Bonferroni adjusted) |
| --- | --- | --- | --- |
| GO:0022904 | Respiratory electron transport chain | 12.95 | 6.0x10<sup>-6</sup> |
| GO:0022900 | Electron transport chain | 11.95 | 1.2x10<sup>-5</sup> |
| GO:0042773 | ATP synthesis coupled electron transport | 12.27 | 0.00017 |
| GO:0006119 | Oxidative phosphorylation | 11.71 | 0.00024 |
| GO:0019646 | Aerobic electron transport chain | 14.46 | 0.00025 |
| GO:0045333 | Cellular respiration | 6.19 | 0.00027 |
| GO:0042775 | Mitochondrial ATP synthesis coupled electron transport | 13.61 | 0.00037 |
| GO:0009060 | Aerobic respiration | 6.49 | 0.00041 |
| GO:0015980 | Energy derivation by oxidation of organic compounds | 4.75 | 0.0049 |
| GO:0006091 | Generation of precursor metabolites and energy | 3.78 | 0.0098 |
| GO:0051238 | Sequestering of metal ion | 101.62 | 0.014 |
| GO:1900779 | Fumiquinazoline C metabolic process | 50.81 | 0.039 |
| GO:1900781 | Fumiquinazoline C biosynthetic process | 50.81 | 0.039 |
| GO:0006120 | Mitochondrial electron transport, NADH to ubiquinone | 50.81 | 0.039 |
| GO:0009821 | Alkaloid biosynthetic process | 7.88 | 0.043 |

## Discussion

The data presented here indicate that *A. fumigatus* HosA is required for normal resistance to intracellular oxidant stress, pulmonary epithelial cell invasion and damage, and virulence during invasive pulmonary infection. When tested in the mouse model of invasive pulmonary aspergillosis, the Δ*hosA* mutant exhibited significantly decreased lethality and caused a reduced host inflammatory response. However, the pulmonary fungal burden of mice infected with the Δ*hosA* mutant was similar to that of mice infected with the Af293 parent strain. We have observed a similar pattern of attenuated lethality with normal to increased pulmonary fungal burden in mice infected with *A. fumigatus* mutants that have defects in the production of secondary metabolites and toxins such as gliotoxin and the Aspf1 ribotoxin (20). Thus, the virulence defect of the Δ*hosA* mutant may be due to a decrease in secondary metabolite or toxin production.

The RNA-seq data indicated that the mRNA levels of genes in the fumitremorgin B and fumigermin biosynthetic pathways were significantly reduced in the HosA mutant. Fumetrimorgin biosynthesis has been found to be controlled by Skn7 (Afu6g12522) and RofA (Afu1g14945) (21). We found that deletion of *hosA* had no effect on *skn7* and *rofA* transcript levels, suggesting that HosA governs expression of the fumetrimorgan gene cluster independently of Skn7 and RofA.

It is unlikely that reduced expression of the fumetrimorgan and fumigermin gene clusters was the cause of the attenuated pathogenicity of the Δ*hosA* mutant. Although fumetrimorgan B is known to be toxic to mice (22), an *A. fumigatus* Δ*fmqA* mutant has wild-type virulence in the zebrafish model of infection (23). Fungermin is known to inhibit germination of *Streptomyces rapamycinicus* during the bacterial-fungal interaction (24). However, its role in *A. fumigatus* virulence has not been determined. More importantly, our previous *in vivo* RNA-seq data indicate that the fumetrimorgan B and fumigermin biosynthetic gene clusters are expressed at a very low level in the lungs of immunosuppressed mice with invasive pulmonary aspergillosis (Table 3) (25). By contrast, the biosynthetic gene cluster for gliotoxin, a secondary metabolite that plays an unequivocal role in *A. fumigatus* virulence (20, 26–28), is expressed at much higher levels. Taken together, these data indicate that although the virulence defect of the Δ*hosA* mutant is likely due to reduced production of one or more secondary metabolites, they are unlikely to be products of the fumetrimorgan or fumigermin gene clusters.

**Table 3.**
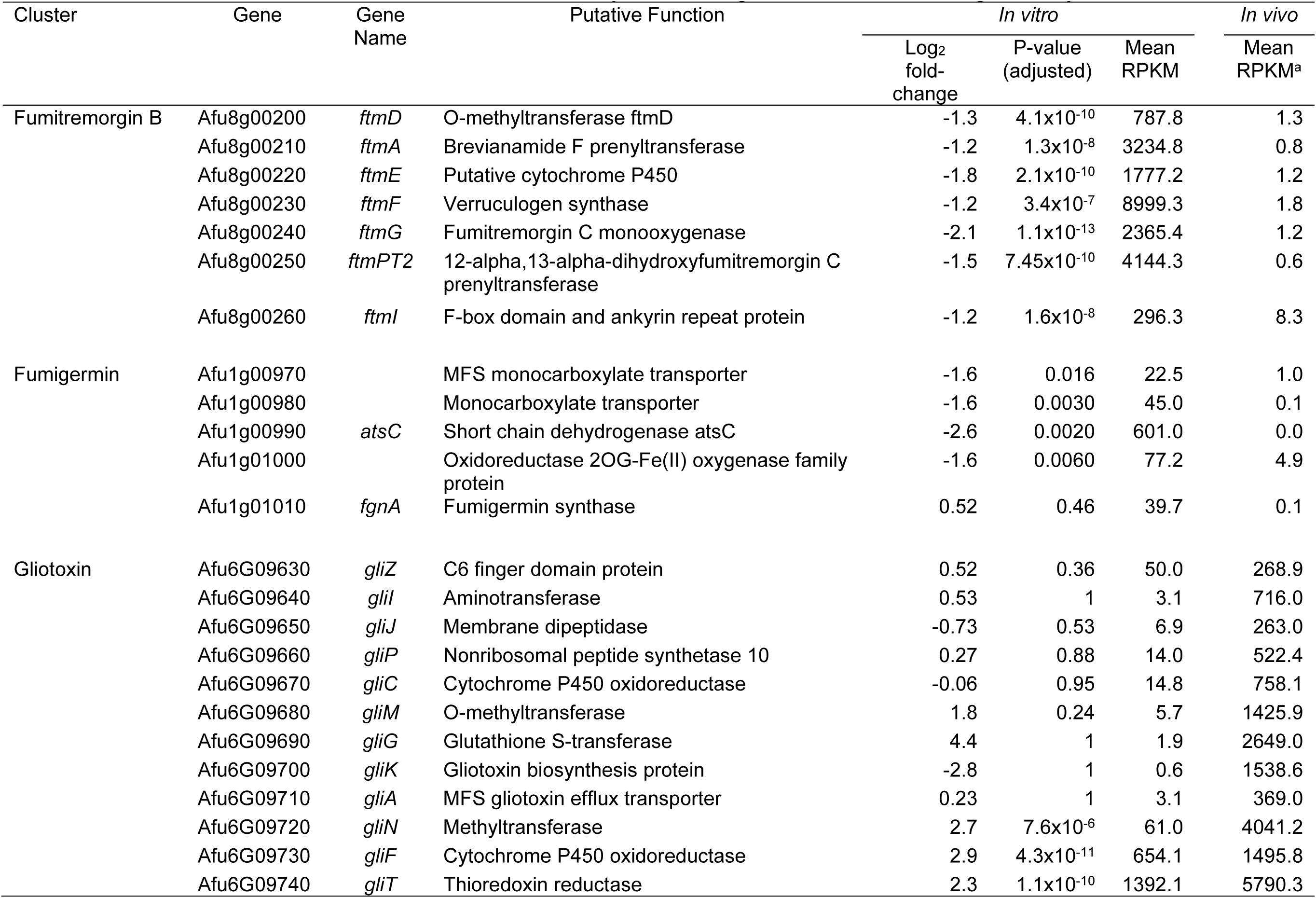
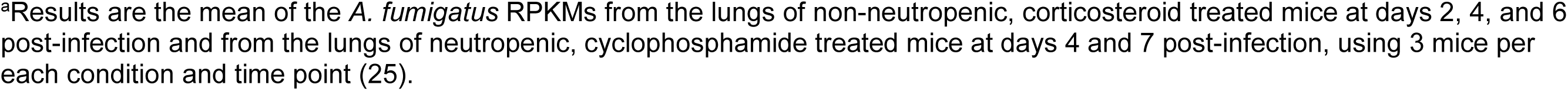
*In vitro* and *in vivo* mRNA levels of selected secondary metabolite gene clusters that are regulated by HosA.

| Cluster | Gene | Gene Name | Putative Function | *In vitro* |  |  | *In vivo* |
| --- | --- | --- | --- | --- | --- | --- | --- |
|  |  |  |  | Log<sub>2</sub> fold-change | P-value (adjusted) | Mean RPKM | Mean RPKM<sup>a</sup> |
| Fumitremorgin B | Afu8g00200 | *ftmD* | O-methyltransferase *ftmD* | -1.3 | 4.1x10<sup>-10</sup> | 787.8 | 1.3 |
|  | Afu8g00210 | *ftmA* | Brevianamide F prenyltransferase | -1.2 | 1.3x10<sup>-8</sup> | 3234.8 | 0.8 |
|  | Afu8g00220 | *ftmE* | Putative cytochrome P450 | -1.8 | 2.1x10<sup>-10</sup> | 1777.2 | 1.2 |
|  | Afu8g00230 | *ftmF* | Verruculogen synthase | -1.2 | 3.4x10<sup>-7</sup> | 8999.3 | 1.8 |
|  | Afu8g00240 | *ftmG* | Fumitremorgin C monooxygenase | -2.1 | 1.1x10<sup>-13</sup> | 2365.4 | 1.2 |
| | Afu8g00250 | *ftmPT2* | 12-α,13-α-dihydroxyfumitremorgin C prenyltransferase | -1.5 | 7.45x10<sup>-10</sup> | 4144.3 | 0.6 |
|  | Afu8g00260 | *ftmI* | F-box domain and ankyrin repeat protein | -1.2 | 1.6x10<sup>-8</sup> | 296.3 | 8.3 |
| Fumigermin | Afu1g00970 |  | MFS monocarboxylate transporter | -1.6 | 0.016 | 22.5 | 1.0 |
|  | Afu1g00980 |  | Monocarboxylate transporter | -1.6 | 0.0030 | 45.0 | 0.1 |
|  | Afu1g00990 | *atsC* | Short chain dehydrogenase *atsC* | -2.6 | 0.0020 | 601.0 | 0.0 |
|  | Afu1g01000 |  | Oxidoreductase 2OG-Fe(II) oxygenase family protein | -1.6 | 0.0060 | 77.2 | 4.9 |
|  | Afu1g01010 | *fgnA* | Fumigermin synthase | 0.52 | 0.46 | 39.7 | 0.1 |
| Gliotoxin | Afu6G09630 | *gliZ* | C6 finger domain protein | 0.52 | 0.36 | 50.0 | 268.9 |
|  | Afu6G09640 | *gliI* | Aminotransferase | 0.53 | 1 | 3.1 | 716.0 |
|  | Afu6G09650 | *gliJ* | Membrane dipeptidase | -0.73 | 0.53 | 6.9 | 263.0 |
|  | Afu6G09660 | *gliP* | Nonribosomal peptide synthetase 10 | 0.27 | 0.88 | 14.0 | 522.4 |
|  | Afu6G09670 | *gliC* | Cytochrome P450 oxidoreductase | -0.06 | 0.95 | 14.8 | 758.1 |
|  | Afu6G09680 | *gliM* | O-methyltransferase | 1.8 | 0.24 | 5.7 | 1425.9 |
|  | Afu6G09690 | *gliG* | Glutathione S-transferase | 4.4 | 1 | 1.9 | 2649.0 |
|  | Afu6G09700 | *gliK* | Gliotoxin biosynthesis protein | -2.8 | 1 | 0.6 | 1538.6 |
|  | Afu6G09710 | *gliA* | MFS gliotoxin efflux transporter | 0.23 | 1 | 3.1 | 369.0 |
|  | Afu6G09720 | *gliN* | Methyltransferase | 2.7 | 7.6x10<sup>-6</sup> | 61.0 | 4041.2 |
|  | Afu6G09730 | *gliF* | Cytochrome P450 oxidoreductase | 2.9 | 4.3x10<sup>-11</sup> | 654.1 | 1495.8 |
|  | Afu6G09740 | *gliT* | Thioredoxin reductase | 2.3 | 1.1x10<sup>-10</sup> | 1392.1 | 5790.3 |

<sup>a</sup>Results are the mean of the *A. fumigatus* RPKMs from the lungs of non-neutropenic, corticosteroid treated mice at days 2, 4, and 6 post-infection and from the lungs of neutropenic, cyclophosphamide treated mice at days 4 and 7 post-infection, using 3 mice per each condition and time point (25).

HDACs are promising therapeutic targets for the treatment of cancers and fungal infections (29, 30). Hos1, the *C. albicans* ortholog of *A. fumigatus* HosA, has been found to be important for governing cell wall synthesis, polyene resistance, and virulence in this fungus. Also, pharmacologic inhibition of Hos1 is therapeutic in the mouse model of disseminated candidiasis and enhances *in vivo* antifungal activity when combined with amphotericin B (31). Based on these results, we speculate that pharmacological inhibition of HosA would be of therapeutic benefit.

## Materials and Methods

### Ethics statement

The mouse studies were approved by the Institutional Animal Care and Use Committee at the Lundquist Institute for Biomedical Innovation at Harbor-UCLA Medical Center. The mice were housed according to experimental group in HEPA-filtered laminar flow cages with unrestricted access to food and water. The vivarium is managed by the Lundquist Institute for Biomedical Innovation at Harbor-UCLA Medical Center in compliance with all polices and regulations of the Office of Laboratory Animal Welfare of the Public Health Service. The facility is fully accredited by the American Association for Laboratory Animal Care.

### Strains and growth conditions

The *A. fumigatus* Af293 strain was purchased from the American Type Culture Collection (ATCC). The Af293 strain that expresses GFP was constructed as previously described (15). All *A. fumigatus* strains were grown on Sabouraud dextrose agar (Difco) at 37°C for 7-10 days prior to use. Conidia were harvested by gently rinsing the agar plates with PBS containing 0.1% Tween 80 (Sigma-Aldrich), after which the conidia were filtered through a 40 µm cell strainer (Corning) to remove the hyphae. The conidia were enumerated with a hemacytometer. To analyze host cell interactions of live germlings, the conidia were pre-germinated in Sabouraud dextrose broth (Difco) at 37°C for 5.5 h.

### Strain construction

To construct the Δ*hosA* (Afu2g03810) deletion mutant, the transient CRISPR-Cas9 gene deletion system was used (32, 33). The Cas9 expression cassette was amplified from plasmid pFC331 (33), using primers Cas9-F and Cas9-R (Table S2). Two 20 bp protospacer sequences were designed in the 5’ and 3’ regions of the target gene coding sequence (Table S2). The sgRNA expression cassette was constructed by overlap fusion PCR. First, two DNA fragments were amplified from plasmid pFC334 (33) using paired primers sgRNA-F, sgRNA-ss-R and sgRNA-R, sgRNA-ss-F (Table S2). Next, the sgRNA expression cassette was amplified by fusion PCR from the two DNA fragments, using primers sgRNA-F and sgRNA-R (Table S2). The hygromycin resistance (HygR) repair template was amplified from plasmid pVG2.2-hph (34) using paired primers Hyg-F and Hyg-R, which had about 50 bp homology with the upstream 5’ protospacer and the downstream 3’ protospacer. *A. fumigatus* protoplasts were transformed with a mixture containing the HygR repair template, the Cas9 cassette, and the two sgRNA cassettes. Hygromycin resistant clones were screened for deletion of the target gene by colony PCR using primers HosA-ScreenF and HosA-ScreenR (Table S2). The positive clones were also confirmed to have no integration of Cas9 or gRNA, using paired primers Cas9RT-F, Cas9RT-R and sgRT-F, sgRT-R (Table S2).

Different strains of *A. fumigatus* that constitutively expressed GFP were constructed for use in epithelial cell invasion assays. Wild-type Af293 and the Δ*hosA* mutant strains were transformed with the plasmid GFP-Phleo and the Δ*hosA+hosA* complemented strain was transformed with plasmid GFP-pPTRI as previously described (15).

To complement the Δ*hosA* mutant, a 3903 bp fragment including the entire *hosA* protein coding sequence was PCR amplified from genomic DNA using primers HosA-Com-F and HosA-Com-R (Table S2). This fragment was cloned into the NotI site of plasmid p402 (35), which was then used to transform the Δ*hosA* mutant. Phleomycin resistant clones were screened for the presence of the *hosA* gene by colony PCR using primers HosA-RT-F and HosA-RT-R (Table S2). The *hosA* transcript level in the various clones was quantified by real-time RT-PCR using primers HosA-Realtime-F and HosA-Realtime-R (Table S2). The clone in which the *hosA* mRNA expression was most similar to the wild-type strain was used in all subsequent experiments.

### Stress assays

To test the susceptibility of the various strains to cell wall, cell membrane and oxidant stress, serial 10-fold dilutions of conidia ranging from 10^5^ to 10^2^ cells in a volume of 5 μl were spotted onto *Aspergillus* minimal medium agar plates supplemented with 15 µM menadione, 4 mM H_2_O_2_, 200 µg/ml Congo red, 300 μg/ml Calcofluor White, and 5 mM protamine (all from Sigma-Aldrich). Susceptibility to voriconazole was determined by adding approximately 100 conidia in 4µl PBS with 0.01% Tween 20 onto Sabouraud dextrose agar plates containing 0.25 µg/ml voriconazole. Susceptibility to hypoxia was determined similarly except that Sabouraud dextrose plates were incubated in and 1% O_2_. Fungal growth was analyzed after incubation at 37°C for 48-60 h.

### Conidia production

To quantify conidial production, 10⁴ conidia were inoculated at the center of 100-mm Petri dishes containing Sabouraud agar. After 5 d incubation at 37°C, the conidia were harvested by gently rinsing the colony surface 100 times with PBS containing 0.1% Tween 80. The conidia were enumerated using a hemacytometer.

### Cell lines

All human cell lines were purchased from the ATCC. A549 cells were cultured in F-12 K medium (ATCC, 30-2004) supplemented with 10% fetal bovine serum (FBS) (Gemini Bio-Products) and 1% streptomycin and penicillin (Irvine Scientific). HSAEC1-KT (HSAE) cells were cultured in SAGM BulletKit medium (Lonza CC-3119 and CC-4124). Both A549 and HSAE cells were grown in 5% CO_2_ at 37°C. The A549 and HSAEC1-KT cell lines were authenticated by the ATCC. All cell lines had no mycoplasma contamination, as determined using MycoAlert^®^ Mycoplasma Detection Kit (Lonza LT07-218).

### Host cell endocytosis assay

The endocytosis of germlings of the different strains by A549 and HSAE cell lines was determined by our previously described differential fluorescence assay (15). Briefly, 2.5 × 10^5^ host cells were grown to confluency on fibronectin coated glass coverslips in a 24-well tissue culture plate. On the day of the experiment, 10^5^ germlings were added to the host cells and incubated for 2.5 h. At the end of the incubation period, the cells were rinsed with PBS in a standardized manner, fixed with 4% paraformaldehyde, and stained with a rabbit polyclonal anti-*Aspergillus* antibody (Meridian Life Science, Inc.) followed by an AlexaFluor 568-labeled goat anti-rabbit antibody (Life Technologies). The coverslips were mounted inverted on microscope slides and viewed under epifluorescence. At least 100 organisms per coverslip were scored for endocytosis and each strain was tested in triplicate in three independent experiments.

### Cell damage assay

The amount of damage to A549 and HSAE cells caused by direct contact with *A. fumigatus* was determined using a ^51^Cr release assay as described previously (16). The inoculum was 5 × 10^5^ conidia per well and the incubation time was 18 h.

### Mouse models of invasive aspergillosis

To assess the virulence of different strains during invasive aspergillosis, we used the triamcinolone immunosuppressed mouse model (18, 19)(18, 19)(18, 19). Briefly, 6-week old male Balb/C mice (Taconic Laboratories) were immunosuppressed with triamcinolone 30 mg/kg administered subcutaneously on days −1 and +1 relative to infection. To prevent bacterial infections, enrofloxacin (Baytril, Western Medical Supply) was added to the drinking water at a final concentration of 0.005% on day −1 relative to infection. The mice were infected by placing them in a chamber into which 1.2×10^10^ conidia were aerosolized. Control mice were immunosuppressed but not infected. The mice were monitored for survival twice daily by an unblinded observer and moribund mice were humanely euthanized. The survival experiments were repeated twice, and the results were combined for a total of 16 mice per group.

To determine the pulmonary fungal burden and cytokine levels, 12 mice per group were immunosuppressed and infected in the aerosol chamber. After 5 d of infection, the mice were sacrificed and their lungs were harvested and homogenized in 1 ml PBS with 10 µl proteinase inhibitor (Sigma-Aldrich, P8340). The homogenates were clarified by centrifugation and the cytokine content of the supernatant was determined using a Luminex Multiplex Assay. The DNA in the pellet was isolated using the ZR Fungal/Bacterial DNA MiniPrep^TM^ kit following the manufacturer’s directions. The fungal DNA content in the various samples was quantified by real-time qPCR using primers ASF1 and ADR1 using the 2^−ΔΔCT^ method with mouse GAPDH-F and GAPDH-R primers as the reference (Table S2).

### RNA-seq analysis

For RNA-seq analysis, 1.5 × 10^8^ conidia of the wild-type and Δ*hosA* strains were added to 300 ml modified Aspergillus minimal medium without added iron and zinc and incubated for 24 h at 37°C in a shaking incubator. The resulting hyphal mat was collected by filtration and the RNA was extracted using the RNeasy Plant Minikit (Qiagen) following the manufacturer’s instructions. The RNA was treated with DNase using the Turbo DNA-free kit (AM1907, Invitrogen). RNA-seq libraries were prepared and 150 bp paired end reads were generated by Genohub Inc. Sequencing reads were mapped to the Af293 reference genome using HISAT2, a short-read aligner. We obtained an average of 40 million aligned reads per sample. Differential expression analysis using raw read counts was done using the DESeq2 R package (36). A gene was considered differentially expressed if the absolute fold-change was ≥ 2.0 and the false discovery rate was ≤ 0.05.

All of the raw sequencing reads from this study are available at the NCBI Sequence Read Archive (SRA) under BioProject accession number PRJNA1361599.

## Acknowledgements

This work was supported in part by NIH grants 5R01AI162802 to SGF and WSM, 2R01AI143198 to WSM, and U19 AI110820 to VMB.

---

**Fig. S1.**
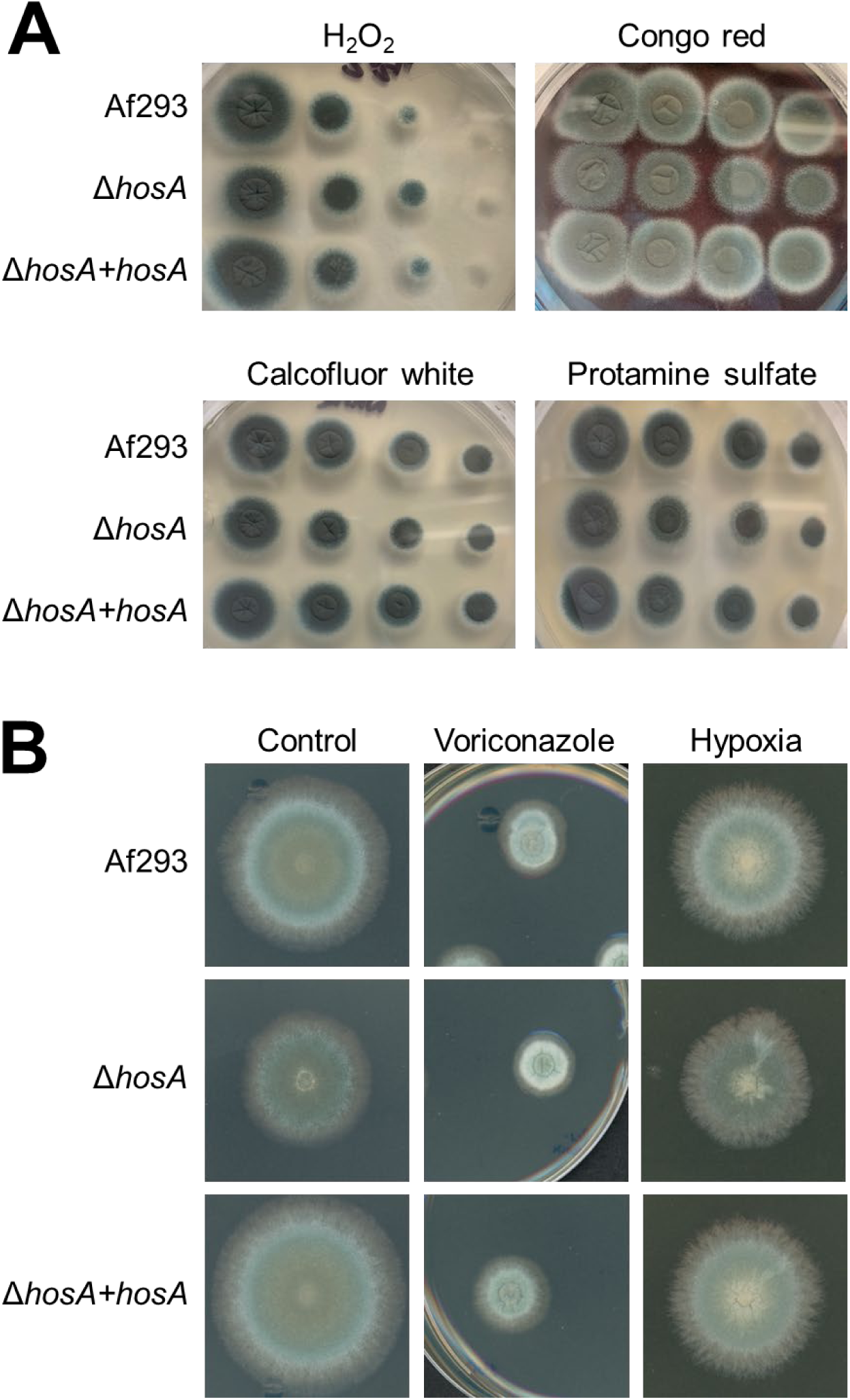
Stress susceptibility of the Δ*hosA* mutant. (A) Susceptibility of the indicated *A. fumigatus* strains to 4 mM H_2_O_2_, 200 µg/ml Congo red, 300 μg/ml Calcofluor white, and 5 mM protamine sulfate. (B) Growth of the indicated strains in the presence of 0.25 µg/ml voriconazole or 1% O_2_. Colonies were imaged after growth at 37°C for 48 h (A) or 60 h (B).

